# Enhancing genetically engineered *Escherichia coli* bioreporters for the detection of buried TNT-based landmines

**DOI:** 10.1101/2023.07.28.550952

**Authors:** Eden Amiel

## Abstract

TNT-based buried landmines are explosive devices consisting of mainly 2,4,6-trinitrotoluene (2,4,6-TNT), but also a 2,4-dinitrotoluene (2,4-DNT) impurity. It is reported that some 2,4-DNT impurity vapors slowly leak through land mine covers and into the soil above them, making this compound a potential indicator for the presence of buried landmines. There are several approaches to identify the presence of chemicals in the environment, one of which is based on a microbial detection system. This approach was previously used to address the challenge of remotely identifying buried landmines by applying genetically engineered microbial whole cell biosensors capable of sensing trace signatures of the explosive compound 2,4,6-TNT and its degradation product, 2,4-DNT. Upon exposure to 2,4-DNT, the genetically engineered bacterial reporter strains, containing a genetic fusion between a sensing element for 2,4-DNT (*yqjF* gene promoter) and a fluorescence/ bioluminescence reporting element (GFP/ *luxCDABE* genes), generate a measurable dose-dependent signal within the whole cell biosensor system. This biological component of a comprehensive remote sensing solution was significantly improved after several rounds of random mutagenesis to the *yqjF* promoter region using a prone-to-error PCR and selection of the best mutants. In the present project further significant improvements were obtained in signal intensity, response time, and in the limit of detection, following an additional round of random mutagenesis. Two new mutants, named A10 and C5, were identified. The new mutants present high induction levels both in terms of Ratio (sample/control) and delta (sample-control), making these strains more amenable to remote detection. To isolate a more sensitive mutant, the 3rd round of random mutagenesis library (about 1000 strains) was re-exposed to lower concentration of 2,4-DNT, but no better candidate was isolated. Possible alternative approaches to lower the detection threshold of the system, such as the use of permeability mutants, are now being tested.

## INTRODUCTION

Landmines are containers of explosive material with detonating systems that are triggered by contact with a person or vehicle, creating an explosive blast designed to hinder or ultimately immobilize and put the advancement of the enemy into a halt [1]. Mine and countermine technologies and techniques have evolved over the past 3,000 years that by the 20th century, the antipersonnel mine grew into a highly effective weapon and combat multiplier.

### History of the weapon

During the First World War the use of battle tanks drove the development of the antitank landmine and later the antipersonnel landmine, designed to prevent enemy soldiers from removing the antitank landmines [2]. Between 1918 and 1939 the development and use of the antipersonnel landmine became a priority among military strategists putting it into use throughout Poland, Russia and Korea. AP’s were then used in World War II to defend military positions and socioeconomic targets, channel the movement of troops to certain routes from which they can’t deviate, and to cause chaos and economic dislocation as political weapons [3]. Subsequently, AP’s were widely deployed around the world in the period from World War II until the treaty banning their use, production, transfer and stockpiling was adopted in 1997 [4]. As a result, dozens of nations have mine-affected land that must be cleared before it can be returned to productive use. Long after wars are over, landmines make land unusable for farming, schools or living, preventing people from rebuilding lives torn apart by conflict [5]. In 2002, the number of landmines planted around the world was estimated to be above 110 million spread in 70 countries, and the time required removing them around 1100 years [6], mainly due to the risks and complications involved in landmines detection and removal.

### Explosives trace signatures and detection techniques

Approximately 80% of all landmines are TNT-based, meaning the main explosive found in them is comprised of 2,4,6-trinitrotoluene (2,4,6-TNT). These also contain additional manufacture impurities and degradations products including 1,3-dinitrobenzene (1,3-DNB) and 2,4-dinitrotoluene (2,4-DNT) [7,8]. Leakage occurs through cracks in the mine casing and vapors diffuse through the plastic housing of the mine. Some of the emitted explosive traces will be adsorbed to soil constituents and the rest will travel away from the mine, migrating to the ground surface in vapor form, thus serving as a potential indicator for buried landmines [7,9,10].

The identification of these trace compounds has served as a basis for miscellaneous approaches for landmine biodetection [11]. This work is based on using the environmentally more stable 2,4-DNT as an indicator for TNT-landmines. Landmines detection methods can be divided into two basic classes: detection of the non-explosive component (mine case or detonator) and detection of the explosive component and its traces. At present, landmine detection is mostly conducted in a manner similar to that employed 60 years ago - based on metal detection and manual characterization of a detected buried object [9,12]. Although its popularity, this method has high false positive rate and it requires immediate proximity to the landmine. Other approaches under development include chemical and physical detection as well as biological detection which are based on the present of trace signatures in the ground above the mine. One of these biological detection techniques is a comprehensive biosensor system.

Biosensors are defined as measurement devices or systems based on the coupling of a biological component with a transducing element, together producing a measurable signal serving as an indicator for the presents of specific chemicals. When compared to analytical methods, biosensors have several advantages including ease of use, low cost, short response times and specificity. All of these combine to allow simplified field applicability [13]. In this study, the biosensor system we used was comprised of whole-cell genetically engineered bacteria assembled by Yagur-Kroll et al. [14] who’s identified in a previous study two *E. coli* endogenous gene promoters: *yqjF* (predicted quinol oxidase subunit) and *ybiJ*, shown to be responsive to 2,4-DNT. These promoters were selected through the scan of 2,000 E. coli reporter strains library, and genetically fused upstream to either fluorescent (GFP) or bioluminescence (*luxCDABE*) reporter genes constructing an *E. coli* whole-cell sensor strain for 2,4-DNT and 2,4,6-TNT. These were then tested for a dose-dependent response to 2,4-DNT in an aqueous solution, in vapors and in soil. All tests indicated that the endogenous promoters are not sensitive enough for landmine detection in situ, and thus improvement of the sensitivity-i.e lowering the detection threshold is required to consider this method of detection as applicable [14]. When testing the system for a response to explosive traces, selection of suitable reporting genes is required. Bioluminescence allows a much faster and more sensitive detection of the target analyte than fluorescence and thus is easier to work with and more efficient when it comes to lab experiments. The main advantage of fluorescence sensing is reporter stability - once induced, the signal keeps accumulating and may be detected for many hours, in some cases even after the cell death [15,16]. The differences between the two systems derives from the fact that bioluminescence is a measure of enzymatic activity (luciferase enzyme), whereas fluorescence quantifies the presence of a protein (green fluorescent protein) [17]. The kind of results that are achieved with the fluorescence system are better suited for the final, in situ detection system, where sufficient time should allow for the signal (a protein) to accumulate and even improve the sensitivity of the detection system. In the same former study [14], the previously constructed *E. coli* whole-cell sensor strain was significantly improved following three rounds of random mutagenesis to the *yqjF* promoter region using a prone-to-error PCR and selection of the best mutants in a ‘directed evolution’ kind of manner. This approach is based on the creation of new variants by random point mutation, screening of the variants with low levels of 2,4-DNT and selecting the improved variants [18]. Following the selection of the improved variants, a dose-dependent experiment is conducted to yet further narrow the search and to select a few of the most enhanced mutants. In this study, further performance improvement of the *yqjF* promoter was achieved in terms of detection threshold, response time and signal intensity through a 4^th^ round of promoter-directed random mutagenesis (RM). Two new mutants, named A10 and C5 were identified, both present high induction levels in terms of Ratio & Delta (see Materials and methods), making these strains more amenable to remote detection.

Each point mutation found in the A10 and C5 strains was than lonely inserted to the promoter region in order to shed some light on the mechanism of action in the induction of yqjF by 2,4-DNT and maybe better understand the possible role of each region in the promoter itself, including possible transcription factors binding sites and other sites important for the transcription of the yqjF gene. Several combinations of numerous point mutations together were also tested for the purpose of learning whether one point mutation can affect the others in a synergetic or an antagonistic kind of way, if at all.

## METHODS AND MATERIALS

### Bacterial strains and plasmids construction

The plasmids and bacterial strains employed and their genotypes are listed in Table 1. Below is a description of the construction of the plasmids generated for the purpose of the present study. *E.coli* K12 strain MG1655 chromosomal DNA was used as a template for promoters amplification in the PCRs. *E.coli* K12 strain DH5α was used as a host for the construction of the plasmids and for the random mutagenesis procedure. *E. coli* K12 strain MG1655 was used as a host for all exposure assays in liquid media.

**Table 1.**
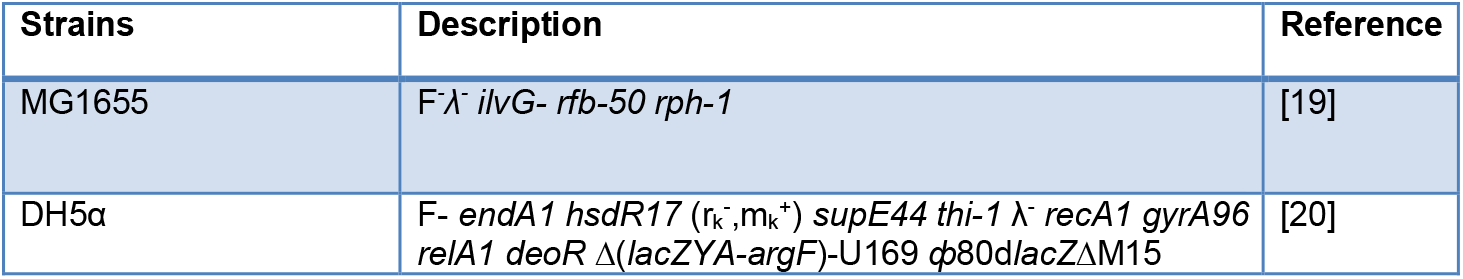
*E.coli* strains used and their genotypes.

The promoter region of *yqjF* was obtained by PCR amplification that introduced *KpnI* and *SacI* restriction sites, seen in Table 2.

**Table 2.**
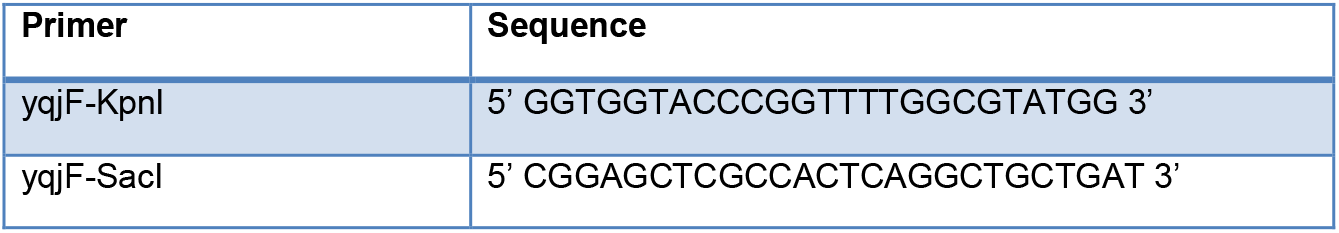
Primers used in this study.

### Random mutagenesis of the *yqjF* promoter region

Another round of random mutagenesis using error-prone PCR was performed to the improved variant *yqjFB2A1#14::lux*, already containing several point mutations in its promoter region (Figure S2), constructed in a previous work following three rounds of random mutagenesis [14]. For the technique to work properly, it was important to use a Taq DNA polymerase which does not have proof-reading ability. Following the error-prone PCR reaction, the PCR products were gel-purified, digested with *KpnI* and *SacI* restriction enzymes [18], and ligated into a promoterless *pBR2TTS:*:*luxC*DABE plasmid (Figure S1), digested with the same restriction enzymes. The products of the ligation reaction were used to transform *E. coli* strain DH5α to generate the variants library. The library was then screened for improved variants as follows: Each colony was inoculated into an individual well of a 96-well microtiter plate, containing 150μl of LB medium supplemented with ampicillin (100μg/ml); two control wells in each plate were occupied by the wild-type strain. This step has been performed by MICROLAB® STAR line (Hamilton, USA) robotic system. The plate was incubated overnight at 37 °C with shaking. Next, 10-μl aliquots of the culture were transferred into two opaque white 96-well microtiter plates with a clear bottom: one containing in each well 90μl of LB supplemented with 25 mg/L 2,4-DNT and the other with 90μl LB supplemented with 0.16 % ethanol. The plates were incubated at 37 °C, and optical density and bioluminescence of the cultures were measured every 20 min using the MICROLAB robotic system’s plate reader. Fluorescence and luminescence values were recorder and displayed as the instrument’s arbitrary relative fluorescence (RFU) and relative luminescence units (RLU), respectively. Blank values were subtracted from the optical density measured, and then the RLU values were divided by the optical density (O.D) of each well. RLU(DNT) and RLU(control) denote the luminescence (after division by the O.D) in wells containing bacteria with or without 2,4-DNT, respectively. The induction ratio for each reporter variant was calculated as follows:

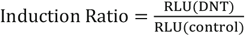

The difference in the signal intensity in the presence and absence of the inducer (ΔRLU) was calculated as follows: ΔRLU = RLU(DNT) − RLU(control) Response sensitivity was determined by calculating the EC_200_ value, denoting the effective inducer concentration causing a twofold increase in the response ratio. Lower EC_200_ values reflect greater sensitivity and a lower detection threshold of the tested chemical by the reporter strain employed [21,22]. Reporter variants that displayed a higher induction ratio, a lower 2,4-DNT detection threshold, or a shorter response time than the wild type were selected for further analysis by exposure to a 2,4-DNT concentration series. The selection procedure of the best variants for further testing was done using an algorithm in MATLAB, as explained below.

### Genetic algorithm as part of a programmed evolutionary solution

A genetic algorithm (GA) script containing multiple functions was developed based on the previous work of Yaara Moskovitz and Tal Elad [14]. The previously programmed algorithm had some limitations that this current GA was able to resolve (see ‘Matlab’ in supplements). The improved GA served us as a search method that mimics the process of directed biological selection. The algorithm operates on a population of potential variants applying the principle of ‘survival of the fittest’ to find better and better candidates suited for our needs. In our fitness-based selection model, the fitness value actually represents the mutant’s ability to detect explosive trace signatures rather than its traditional definition used in several other models [23]. For this, a customized function that assigns fitness values for each variant was developed. At first, the script automatically calculates the induction ratio and the *ΔRLU* for the first 180 minutes of measurements and creates two kinetic graphs for each one of the variants: one for the ratio and one for the *ΔRLU* (Figure S6). Each graph is represented by a function equation - *f(x)*. Next, the fitness values/ranking (*FitnV*) of each variant is calculated by the equation: 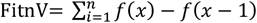 that is the sum of the differences between adjacent elements of X along the function *f(x)*. This value is basically a number that describes how much the *RLU* per OD changed over time and is a good estimation for the mutant’s ability to detect landmines. For fitness-based competition, the FitnV values were organized into a list in descending order. The first 10 values of the two lists (ratio & Δ*RLU*) were intersected, and the variants with the highest FitnV value in both lists were selected for further analysis.

### Exposure to a 2,4-DNT concentration series

The bacterial reporter strains were grown overnight at 37°C in LB medium with 100μg/ml ampicillin. Then, the cells were diluted 100-fold in fresh medium, and regrown with shaking at 37°C to an early exponential growth phase (O.D_600_ ≈ 0.2). In the next step an opaque white 96-well microtiter plates have been filed, so that each well contained 50μl of a predetermined concentration of 2,4-DNT dissolved in 200 mM Tris-base pH 7 and 5μl of x10 concentrated LB media. Into each well, culture aliquots (45μl) were transferred. Luminescence was measured at 37°C every 10 minutes using a VICTOR^2^ plate reader (Wallac, Turku, Finland). All experiments were carried out in duplicate, and were repeated at least 3 times. To assess the activity of the tested reporter strains, *ΔRLU* and induction ratio were calculated as explained above.

### Single and multiple-point mutations

For the study of the effect of specific single nucleotide alterations, primers harboring the point mutations (PM) of interest were used to perform a standard PCR, inserting PM’s to specific areas in the process. All insertions were confirmed by sequencing.

## RESULTS

It was shown in a previous study that sensor performance measured by signal intensity, time of detection, and/or detection threshold, could be improved by a directed evolution procedure based on the insertion of random mutations in the promoter element of the sensor [18]. This procedure has been applied to the *yqjF* promoter prior to this study, performing three rounds of random mutagenesis (RM) PCR and ultimately generating an enhanced sensor called *yqjFB2A1#14::lux* [14]. In the current study, a 4^th^ round of RM was performed using *yqjFB2A1#14::lux* as the template for the error-prone PCR procedure, in order to further improve this sensor’s performance. Following the procedure, several candidates were chosen for an exposure test to a 2,4-DNT concentration series. The best performers obtained from the 4^th^ round RM followed by an exposure to a 2,4-DNT concentration series were the variants *yqjF4th-****A10****::lux & yqjF4th-****C5****::lux* (Figure 1).

**Figure 1.**
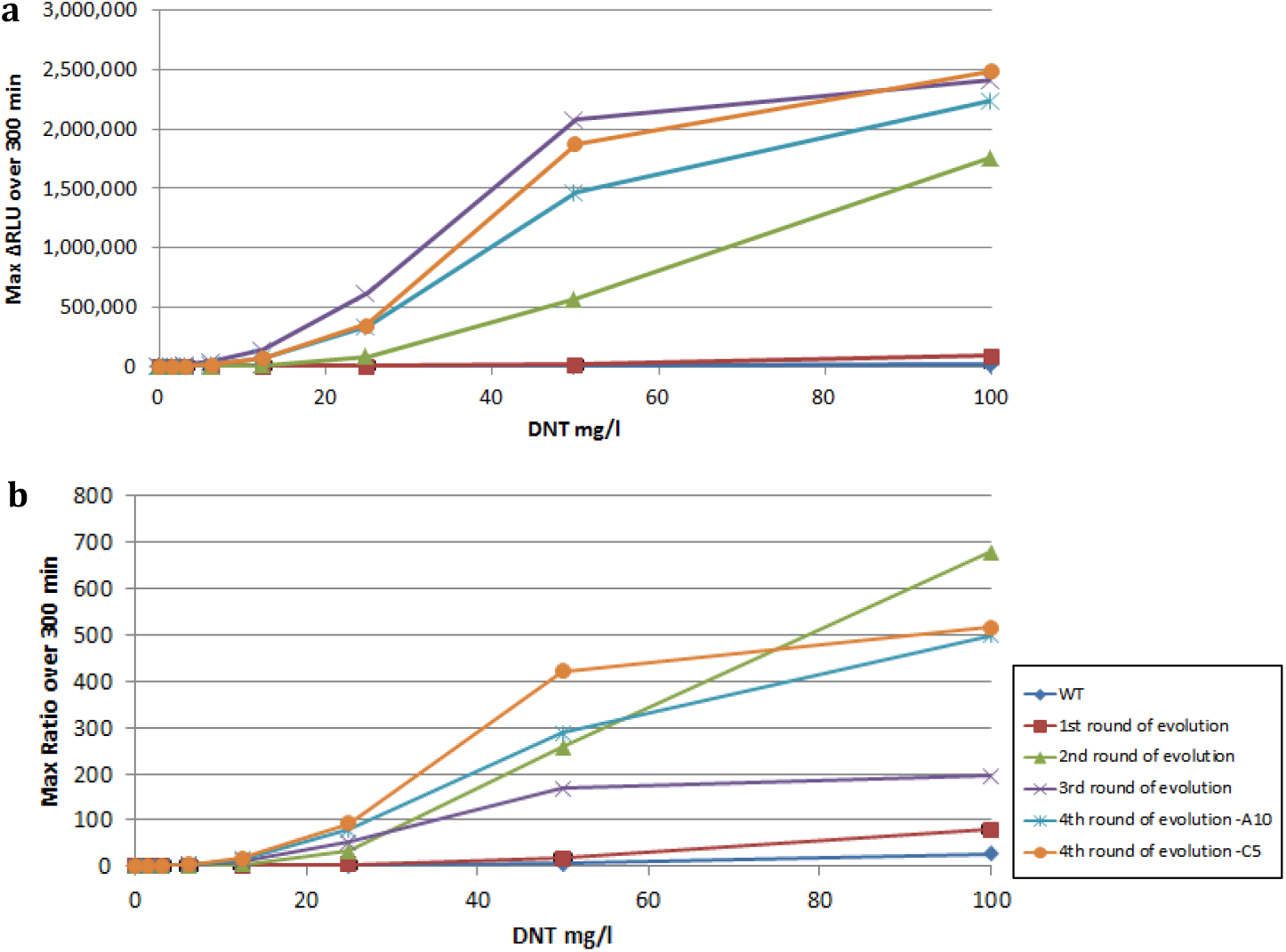
Induction of the original *yqjFwt* and improved variants by 2,4-DNT: dose-dependent response. **a** increase in luminescence intensity (ΔRLU); **b** Response ratio. The data presented are maximal values obtained in the course of a 300min exposure.

For **A10**, the response ratio value was up to 42-fold higher than the wild type (287 as compared to 6.8 for 50 mg/L 2,4-DNT), the *ΔRLU* values were over 400-fold higher (1,167,694 as compared to 2,795 for 50 mg/L 2,4-DNT). For **C5**, the response ratio value was up to 62-fold higher than the wild type (421 as compared to 6.8 for 50 mg/L 2,4-DNT), the *ΔRLU* values were over 500-fold higher (1,441,933 as compared to 2,795 for 50 mg/L DNT).

Furthermore, the response time for 25mg/L 2,4-DNT (Figure 3), defined as the time the response ratio reaches a value of 2 in this concentration, was shortened from about 100 minutes in the WT to approximately 70 minutes in both *A10 & C5*, and as opposed to 80 minutes in the former *yqjFB2A1#14* mutant (Table S1). The response sensitivity/detection threshold indicated by the EC_200_ was considerably improved as well, with EC_200_ values reaching as low as 4.1mg/L of 2,4-DNT (Figure 2). Sequencing and comparing the promoter regions of the two improved variants revealed that *yqjF4th-A10::lux* has an addition of one point mutation compared to the former *yqjFB2A1#14::lux* and *yqjF4th-A10::lux* has four additional ones (Figure S2).

**Figure 2.**
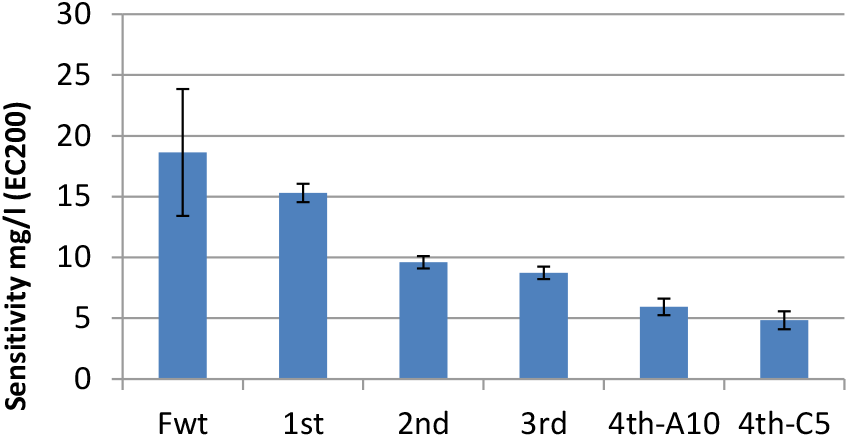
Detection threshold, expressed as EC_200_- the sample concentration yielding a ratio of 2 (measured over 120 minutes).

**Figure 3.**
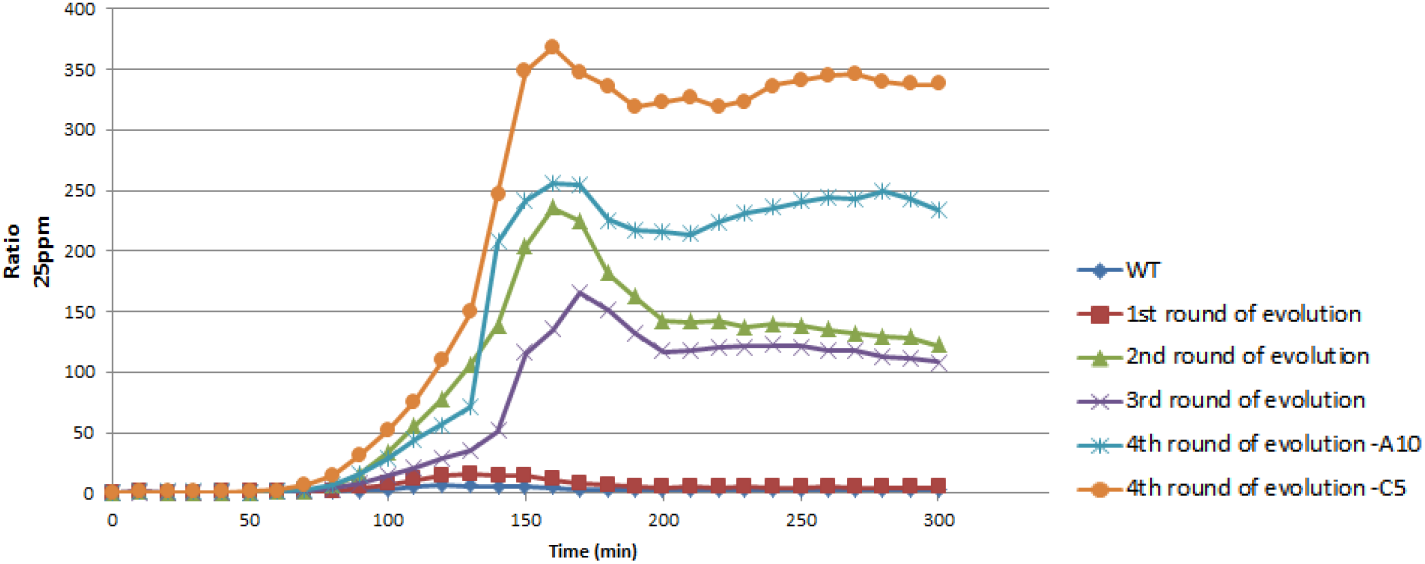
Response ratio of the original *yqjFwt* and improved variants in liquid phase over 300 minutes, calculated at 25 mg/L of 2,4-DNT.

Using different bioinformatics data bases such as RegulonDB [24] and Ecocyc [25] it was shown in a previous study [14] that the promoter *yqjF* contains three predicted transcription units, two of which bind the RNA polymerase sigma 32 factor (*yqjFp3 and yqjFp4*) and one binds RNA polymerase sigma 70 factor (*yqjFp2*). Of the new muta-Fwt 1st 2nd 3rd 4th-A10 4th-C5 tions originated in this study, only one found in C5 was located in a predicted regulatory element of the -10 region of σ^32^-dependent *yqjFp4*.

### Point Mutations – Individual & multiple effects

To better understand the effects of each nucleotide alteration that was made in each round of the RM process and for the purpose of using this knowledge to perceptively engineer an even more enhanced sensor, individual and multiple combinations effects of all point mutations were tested.

The dose-dependent max response ratio over 240 minutes of all 14 point mutations that were inserted and tested individually can be seen in Figure S3, while the response ratio at 100mg/L 2,4-DNT of a selected few is also presented (Figure 4). In the same sense, the dose-dependent max *ΔRLU* over 240 minutes of all combinations made between numbers of PMs is shown in Figure S3. The data of a selected few necessary for the discussion are presented as well (Figure 5). Although the *ΔRLU* of the mutant *MG/F124-F133* reached peaked high values of 4×10^6^, it presented a very low response ratio of about 2, possibly due to its high background values – 250 times higher than that of the WT (Figure S4). In all tests the mutant *MG/F138* lowered both the *ΔRLU* and the response ratio values, but when calculating the EC_200_ it showed an increase capability regarding the sensitivity of the sensor (Figure 6).

**Figure 4.**
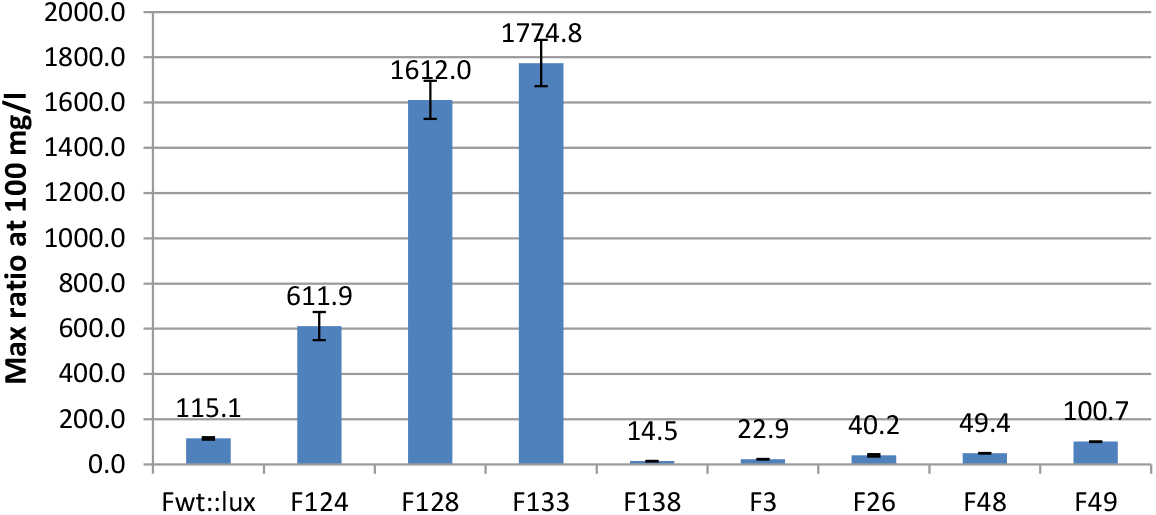
Response ratio of the original *yqjFwt* and selected individual point mutations depict from various strains in liquid phase calculated at a concentration of 100 mg/L 2,4-DNT. Data is max value over 240 minutes.

**Figure 5.**
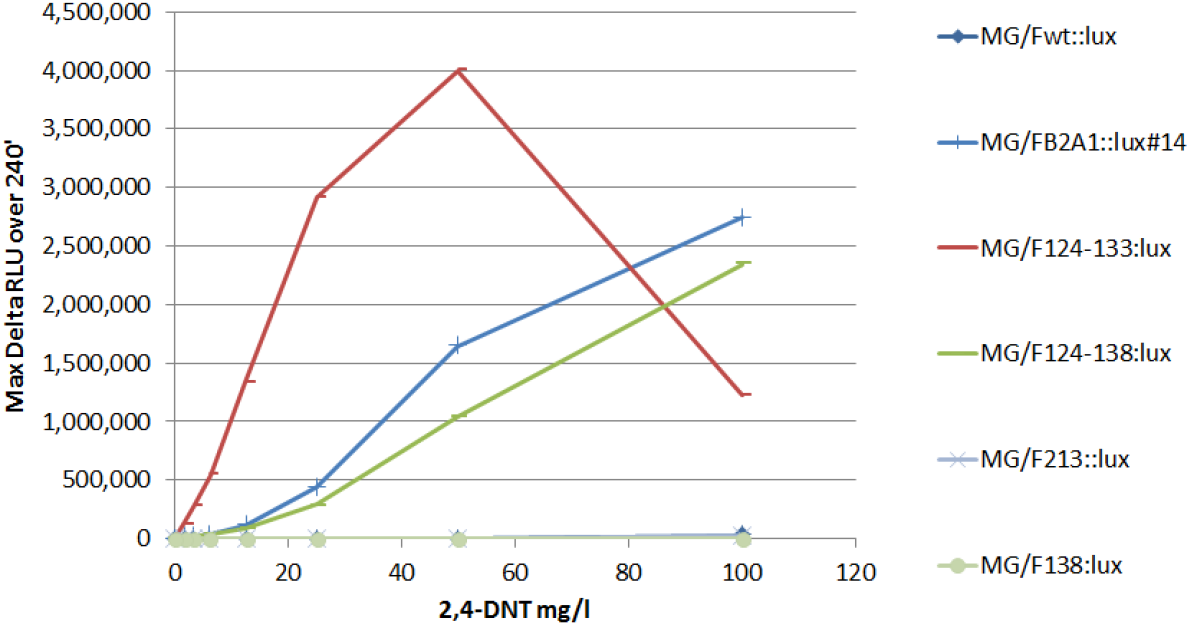
Increase in luminescence intensity (ΔRLU). Dose-depndent response of the original *yqjFwt* and selected individual point mutations and their combinations depicted from various strains in liquid phase, maximal values obtained over 240min.

**Figure 6.**
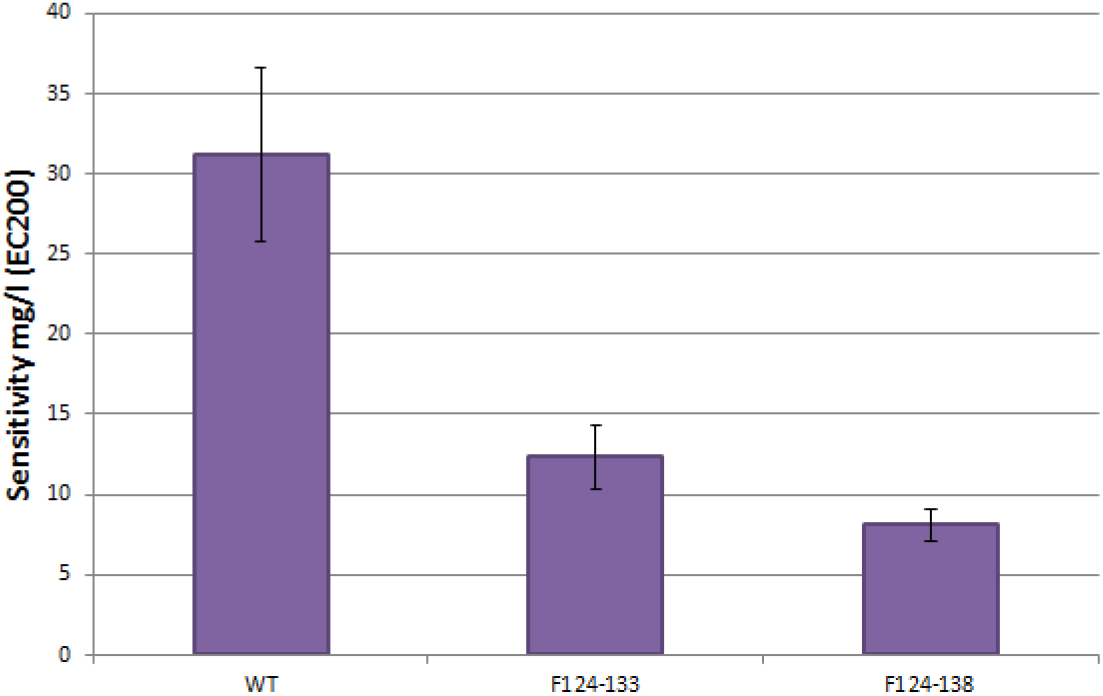
Detection threshold, expressed as EC_200_- the sample concentration yielding a ratio of 2 (measured over 120 minutes). The original *yqjFwt* and selected combinations of several individual point mutations taken from different strains in liquid phase.

## DISCUSSION

Directed evolution by means of random mutagenesis was conducted in this study as one of several solutions proposed in a former study [14] to possibly further enhance the sensor’s performance for the detection of buried TNT-based land mines. Despite the fact that throughout the course of this study we focused our efforts on the *E.coli* reporter strain harboring the *pBR2TTS*:*yqjF::luxCDABE* plasmid, transformation of the detection system from luminescence *luxCDABE-*based to a fluorescence GFP-based for the purpose of field testing was performed and experimented in the lab as well (Results are not shown). In addition, to isolate a more sensitive mutant the 3rd round of random mutagenesis library (about 1000 strains) was re-exposed to lower concentration of 2,4-DNT and scanned with the improved algorithm we have developed, but no better candidate was found. From the 4^th^ round of RM two new mutants - A10 & C5, were identified; both present high induction levels in terms of Ratio (sample/control) and delta (sample-control). Further significant improvements were obtained in signal intensity, response time and the limit of detection as well, making these strains more amenable to remote detection of land mines. However, the limits of detection described (Figure 2) were still above the extremely low vapor concentrations (in the parts per trillion range) of the trace compounds reported to exist above buried land mines [9,26].

When comparing the max *ΔRLU* received over 300 minutes to the corresponding max response ratio it can be observed (Figure 1) that the two new mutants are superior to the last *yqjFB2A1#14* mutant in terms of ratio but not in terms of delta. It can be assumed that this result is not necessarily bad due to mutant’s *#14* high background values (Figure S4) which, in turn, contributes to its relatively high delta values and low ratio values (Figure 3). This suggests that by lowering the background values while keeping the induction signal intensity high as seen in the new mutants, it might be easier to detect the trace signatures found above buried landmines in the field, due to a higher response ratio. Of the four new PMs inserted by the 4^th^ RM in the *C5* strain, one PM named F3 was located in a predicted regulatory element of the prinbow box (−10 region) of σ^32^-dependent *yqjFp4* (Figure S2). When tested the individual effect of this mutant it showed a decreased response than that of the WT in both terms of *ΔRLU* and EC_200_. Three other PMs (F124, F128, F133) displaying promising results, along with others, were introduced in the former 3 rounds of RM done prior to this work [14]. These mutants were later found to be located in the -35 region of the σ^70^-dependent *yqjFp2* transcription unit. In the current study, each of these three PMs was tested individually, every single one presenting a vast enhancement in its response by all aspects. Other results from the individual (Figure 4), and multiple (Figure 5) point mutations tests suggests that although mutant F138 had a negative effect on the performance of the sensor in terms of delta and ratio it seemed to have increased the sensitivity of the sensor when combined with the promising F124-F133 mutant, consisting 3 PMs in the -35 region as explained above (Figure 6). This result is the possible outcome of the reduction in the background values displayed by the F138 mutant (Figure S4) and can serve as a basis for a new approach to further increase the sensor’s sensitivity – by lowering the background values of the strain. In addition, it was surprising to see that the only PM inserted in the *A10* strain (F49) had little or no effect when inserted individually (Figure 4), but when combined with the former PMs found in *yqjFB2A1#14* mutant it had an enhancing response to the former one. This suggests that a specific set of multiple mutations can have a synergetic affect than that of each one individually.

It was concluded in an earlier study conducted by Yaara Moskovitz [27] that due to the dramatically improved sensor’s response results achieved when mutagenesis was applied to the -35 region of the σ^70^-dependent *yqjFp2* transcription unit, sigma 70 plays a major role in the transcription of this gene and that the possible outcome of the new PMs introduced in mutant F124-F133 could be an increment in the affinity of the RNA polymerase-σ^70^ holoenzyme to the *yqjF* promoter. However, based on the results acquired in this study I suggest that a different possible mechanism of action for the induction of the *yqjF* promoter by 2,4-DNT takes place. It might be possible that the induction of this promoter occurs through the σ^32^-dependent *yqjFp4* transcription unit and not the σ^70^-dependent *yqjFp2* transcription unit as proposed before. This conclusion is based on the following observations and assumptions: **(1)** The F3 mutant which introduced a nucleotide alteration in the prinbow box (−10 region) of σ^32^-dependent *yqjFp4* resulted in a decreased activity of the promoter when compared to the WT. **(2)** Sigma factor subunit binds transiently to the core RNA polymerase enzyme and directs it to specific binding sites. Because specific sigma factors are needed at specific promoters, the use of sigma factors gives *E. coli* a powerful mechanism to regulate a group of genes involved in a specific function, such as the heat shock (HS) response genes. RNA polymerase binds to the promoters of these genes only when σ^70^ is replaced with the σ^32^ subunit (specific for heat shock promoters). Which different σ subunits will bind the core enzyme is determined by several factors including its relative cellular levels compared to other σ subunits levels [29]. During normal exponential growth, σ^70^ accounts for 60-95% of the total pool of cellular sigma factors with approximately 700 molecules per cell while σ^32^ accounts for less than 10 molecules per cell [28,29]. *In vitro* experiments at low temperatures have shown that σ^32^ and σ^70^ have similar affinities for RNA polymerase-core binding [30]. This fact suggests that σ^32^ competes with σ^70^ for RNA polymerase binding and thus lowering the levels of free σ^70^ will provide σ^32^ a competitive advantage. To my understanding, this can be the case in our research in which by making mutation in the -35 region of the σ^70^ – dependent transcript we’re not increasing the σ^70^-RNAP holoenzyme’s affinity to the promoter but rather decreasing its affinity. What this does is that it leaves the σ^70^ subunit bound to the core RNA polymerase rendering it unavailable and not allowing it to dissociate, as initiation of transcription is required for the release of the sigma factor. Consequently, the cellular levels of free σ^70^ should drop, increasing the relative cellular levels of σ^32^. This should initiate the transcription of σ^32^ – dependent transcripts, as explained above. In this way, the results we observed could be the result of down-regulating a specific transcript, which in turn up-regulates the transcription of a different, possibly HS related transcript. This explanation can also give us an answer as to why an endogenous *E. coli* promoter is strongly and specifically induced by synthetic chemicals that are not usually found in its environment. A tolerance mechanism against nitrogen compounds and exploitation of the nitro groups as a nitrogen source or the aromatic ring as a carbon source were previously suggested [1]. It can now be assumed that this is true in the case of cell’s exposure to various environmental insults, such as nitrogen or carbon starvation which can induce σ^32^ [31] that will subsequently bind RNAP and initiate the transcription of the *yqjF* gene. The ultimate response of the cell for this case is scavenging: when a particular nutrient becomes limiting, *E. coli* increases the production of proteins that forage for the limiting nutrient [33]. One of these scavenging proteins might include the *yqjF* gene which translates to an inner membrane protein with four predicted transmembrane domains. In addition, it was shown that *E. coli* is able to use 2,4,6-TNT as the sole nitrogen source [32], making this proposed explanation more probable. Although this might be the case, further research is clearly required in order to understand the role of *yqjF* in the metabolism of TNT and 2,4-DNT.

Determination of the relative cellular levels of σ^32^ by means of Western blotting should be considered as a method to further research or to disprove this hypothesis.

In conclusion, by applying the directed evolution approach, further improvements were made towards an operative TNT-based landmine detection system. The new variants displayed higher signal intensity, responded faster and were able to detect lower 2,4-DNT concentrations. Though not yet sufficiently sensitive to detect the low levels reported to exist above buried land mines, the efficiency of the directed evolution technique on the promoter region demonstrated in this work is expected to get us closer to a fully functioning landmine detection system.

Future research endeavors should focus on increasing the sensitivity of the system, possibly by the use of mutant strains of several membrane proteins as a host for the genetic fusion to ultimately lower the efflux of 2,4-DNT from the cell. Another approach is to increase the influx levels of the substance through the fusion of the integral membrane protein OmpF porin gene to an IPTG-inducible lacZ gene promoter. Moreover, to further test the effects of multiple mutations individually inserted, a better understanding of the possible role each mutant acquires is necessary due to the high possible combinations that can be made and tested 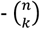.

## SUPPLEMENTS

**Figure S1.**
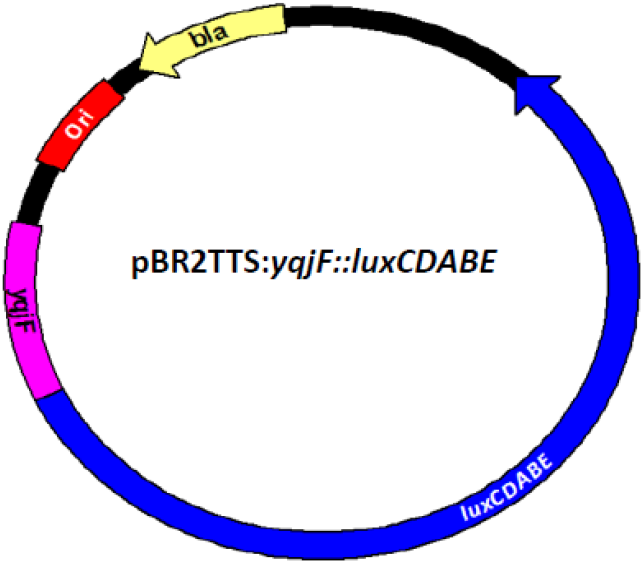
The schematic structure of plasmid pBR2TTS:*yqjF::lux*CDABE, harboring the *yqjF* promoter fused to the *lux* operon, taken from Dr. Sharon Yagur-Kroll’s work [14].

**Figure S2.**
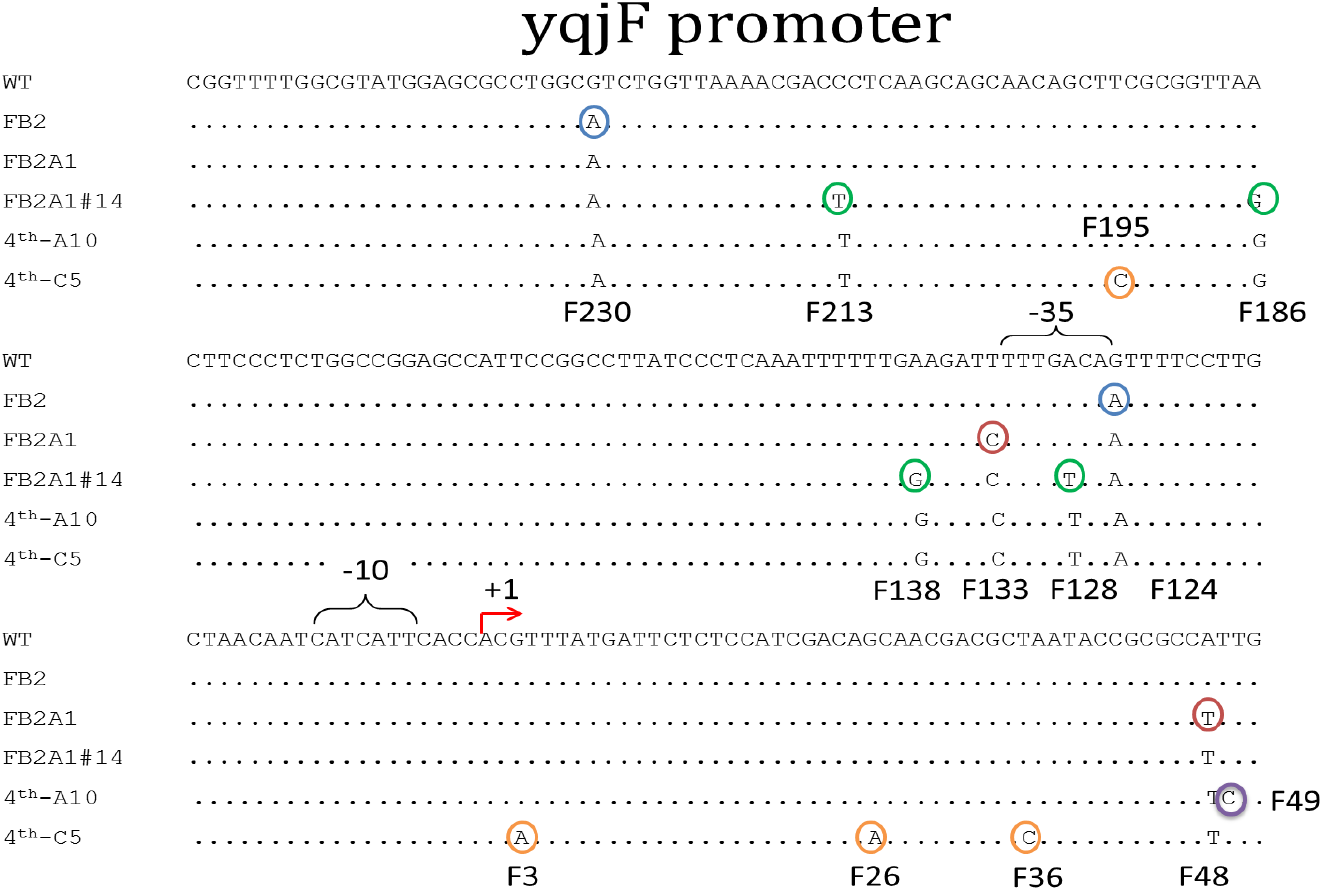
Multiple sequence alignment constructed using two bioinformatics tools Emma and Showalign from the EMBOSS suite. The multiple sequence alignment consists of the promoter sequences of *yqjF*wt, *yqjFB2, yqjFB2A1, yqjFB2A1#14, yqjF14-A10* and *yqjF14-C5*, the variants selected sequential rounds of mutagenesis, respectively. The mutations inserted in the first round are circled in blue, the mutations add in the second round are circled in red, the mutations added in the third round are circled in green, the only mutation added in the 4^th^-A10 is circled in purple and the mutations added the 4^th^-C5 are circled in orange.

### Matlab

In the third round the improved variant picking was based on calculation done by a MATLB script that automatically calculated the induction ratio and the ΔRLU for the first ten measurements (200 minutes). Then it found the coefficients of a polynomial curve of degree one that fit the data for each series of values, ratio and ΔRLU, to the time in a least squares sense. For each series of values the curves slopes were organized into a list in descended order. The first 20 values of the two lists were intersected, and the variants with the highest slope value in both were selected for further analysis. Also, five variants displaying the highest slope values in the ΔRLU list that were not selected before, were selected for further analysis. The problem with this method is in a case where calculating the induction delta measured by RLU(DNT)-RLU(control) and the RLU(control) values are higher than that of the RLU(DNT). In this situation the induction delta received is a negative one and the curve’s slope of the graph that is being calculated by the algorithm doesn’t reflect the real capability of the mutant to be induced by 2,4-DNT. Thus, 2 graphs that will reach the same RLU values but one of them has a negative delta RLU in the beginning of the reaction will result in different ranking, as illustrated below:

**Figure S5.**
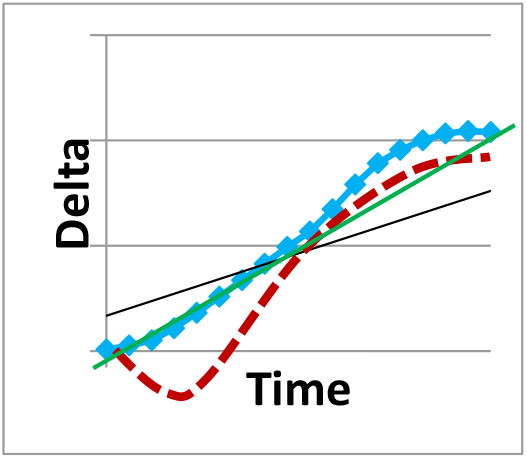
An illustration of the problem the old algorithm had. As seen, the two graphs (displayed in blue and red) have approximately the same delta reaction but because the red graph has negative values of delta in the beginning of the reaction than the slope that is calculated by the algorithm (black line) is smaller than that of the blue graph (green line). This means that the algorithm will give us an output to pick the mutant corresponding to the blue graph and ignore the mutant corresponding to the red graph due to its lower slope, while the two mutants act the same, and the red one might even just be better.

**Figure S6.**
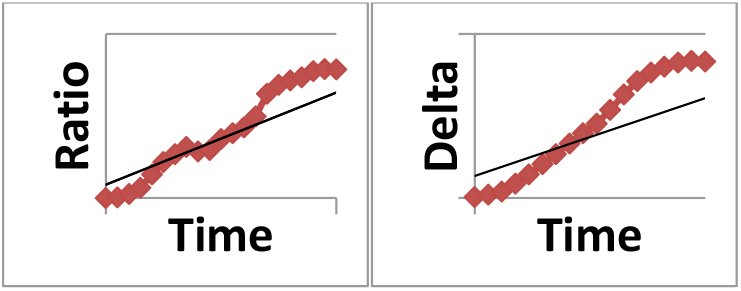
Sample of kinetic graphs build by the Matlab script and the slope that is being taken into account when ranking the best mutants

**Table S1.** Responses of original biorepoter, yqjFwt, and the improved variants to 2,4 -DNT. The response is characterized by three parameters: Signal intensity, expressed as maximum response ratio and maximum ΔRLU in the course of 300min induction (Measured with 25 mg/L 2,4-DNT).Response time which is defined as the time the response ratio reached a value of 2(Measured with 25mg/L 2,4 DNT).Detection threshold, expressed as EC200, the sample concentration yielding a response ratio of 2 (Measured over 120 minutes).

|  | *yqjFwt* | *yqjFB2* | *yqjFB2A1* | *yqjFB2A1#14* | *yqjF4th-A10* | *yqjF4th-C5* |
| --- | --- | --- | --- | --- | --- | --- |
| **Signal intensity (Ratio)** | 2.7 | 3.6 | 33.6 | 53.4 | 80.6 | 93.3 |
| **Signal intensity (ΔRLU)** | 608.5 | 1,983 | 60,673 | 553,895 | 276,049 | 272,957 |
| **Response time (min)** | 100 | 90 | 80-90 | 80 | 70-80 | 60-70 |
| **Detection threshold (EC<sub>200</sub>)** | 18.6±5.2 | 15.3±0.8 | 9.6±0.5 | 8.7±0.5 | 5.9±0.7 | 4.8±0.7 |

**Figure S3.**
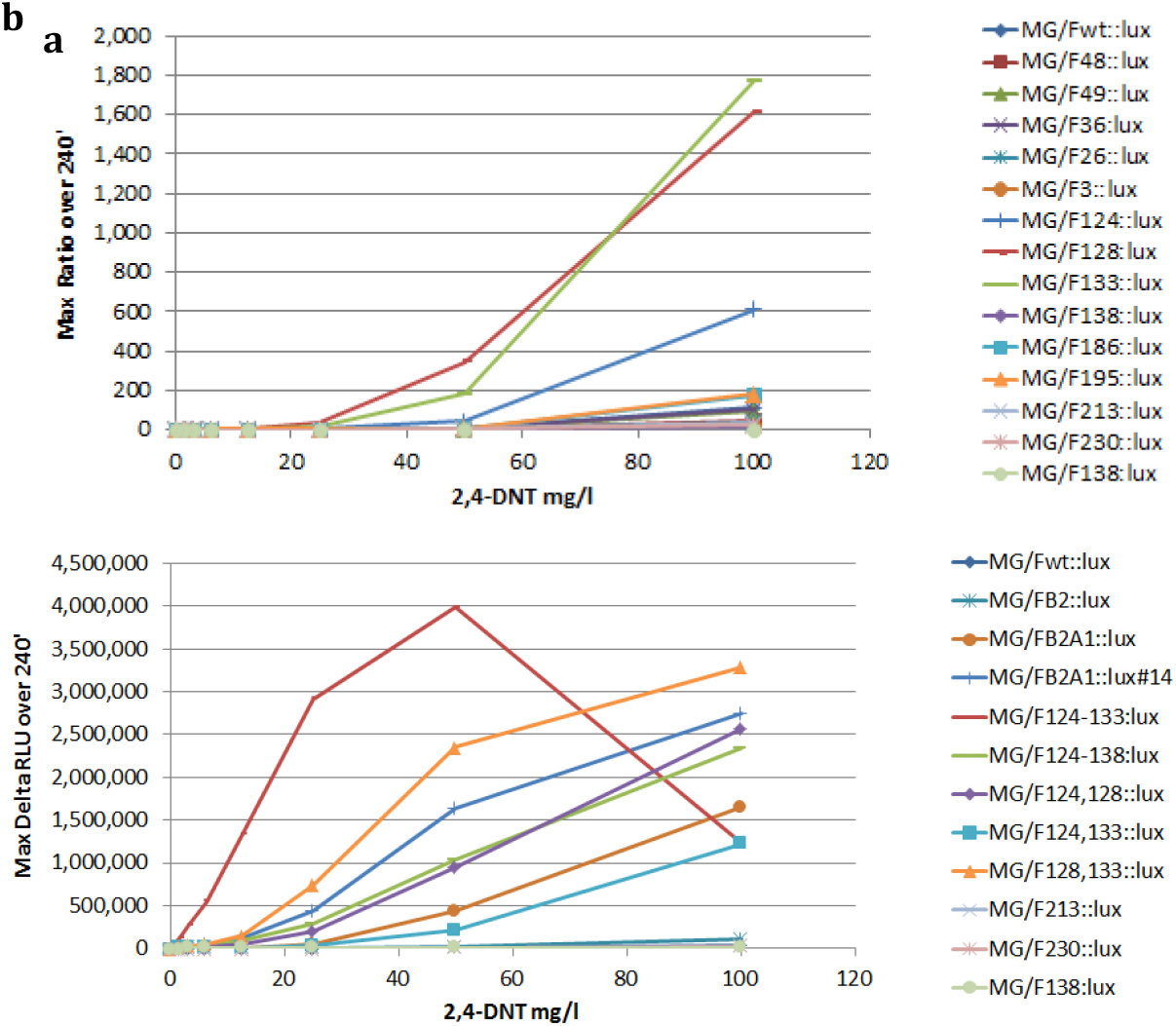
Induction of the original *yqjFwt* and improved variants by 2,4-DNT: dose-dependent response. **a**. Ratio of all 14 point mutations inserted individually **b**. delta RLU of single, and multiple point mutations. All data is displayed is a maximum value over 240min.

**Figure S4.**
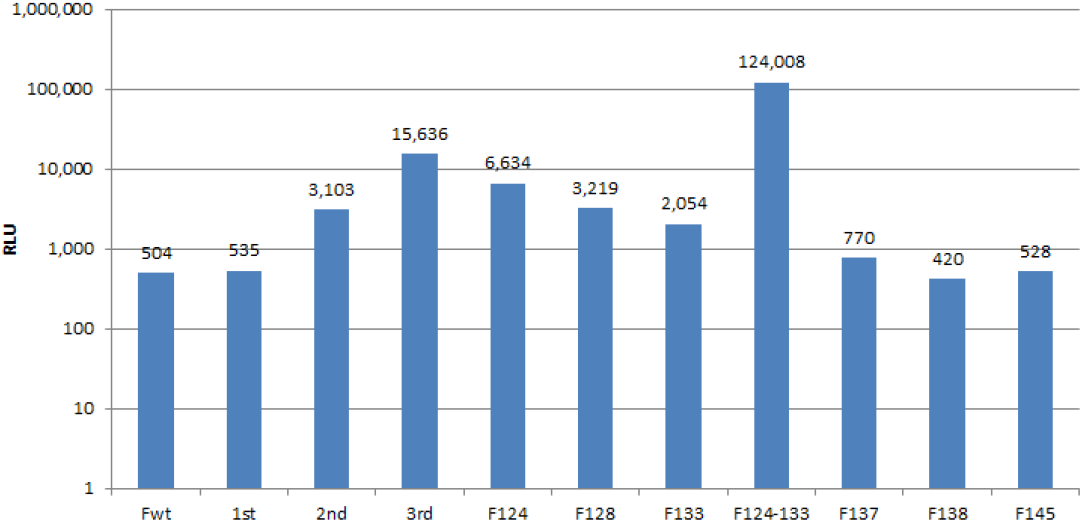
Background RLU values (no DNT present) of the original *yqjFwt* and selected individual point mutations and their combinations depicted from various strains in liquid phase.

